# Flow-driven construction of capillary-scale vessels with predefined geometries in natural hydrogels

**DOI:** 10.1101/2025.06.19.660502

**Authors:** Akira Ono, Ryohei Ouchi, Yusei Kumagai, Masahiko Suda, Tadahiro Yamashita, Yoshihiro Taguchi, Ryo Sudo

**Affiliations:** School of Integrated Design Engineering, Graduate School of Science and Technology, Keio University, Japan; Department of System Design Engineering, Keio University, Japan

**Keywords:** capillary, collagen gel, EndMT, fibrin gel, microvessel, multiphoton lithography

## Abstract

Forming capillary networks with predefined geometries is a critical challenge in engineering complex three-dimensional tissues *in vitro*. While bioprinting and microfluidic technologies have enabled vascular tissue fabrication, precise control over capillary-scale vascularization remains limited. In this study, we investigated vascular formation process along hydrogel microchannels to elucidate mechanisms governing capillary-scale lumen formation. Microchannels were fabricated by femtosecond laser ablation in collagen and fibrin hydrogels. We optimized multiphoton lithography parameters to fabricate microchannels within these hydrogels and analyzed vascular formation along the channels. Luminal vascular structures formed readily in 50-μm channels, while vascular formation failed in 20-μm channels under static conditions, suggesting a significant shift in endothelial organization at the cellular scale. Flow stimulation significantly promoted vessel formation through collective endothelial cell migration and adhesion, whereas static conditions induced endothelial-to-mesenchymal transition. These findings provide key insights into capillary-scale vascularization and contribute to the development of more complex architectures with predefined shapes.

## Introduction

Advances in tissue engineering and stem cell biology have enabled the formation of various three-dimensional (3D) tissues, such as organoids and 3D culture models using microfluidic devices(1–9). However, the morphology of these 3D architectures cannot be precisely controlled, as their morphogenesis predominantly relies on the self-organization of cells. Overcoming this limitation is a critical challenge in engineering complex 3D tissues *in vitro*, particularly in achieving predefined geometries. In particular, the construction of capillaries is crucial, as vital organs depend on capillary networks to facilitate metabolic exchange by delivering oxygen and nutrients while removing metabolic waste within 3D structures.

The *in vitro* construction of capillaries has been achieved using angiogenesis- and vasculogenesis-based models (10–15). Recent advances in microfluidic technology have allowed precise modulation of vascular formation by controlling biochemical and biomechanical stimuli within microfluidic devices (16, 17). Among these stimuli, flow dynamics has been identified as a key regulator of vascular endothelial cell behavior (18–21). Furthermore, co-culture systems incorporating supporting cells, such as pericytes, mesenchymal stem cells, and fibroblasts, have been utilized to promote stable and functional vascular formation (22–24). However, in these models, vascular geometry is predominantly determined by the spontaneous organization of cells, significantly limiting the precise control of vascular structure.

*In vivo*, capillaries exchange substances with parenchymal cells via diffusion, which requires the distance between capillaries and parenchymal tissues to remain within 200 µm to ensure efficient transport (25). Moreover, each organ exhibits a distinct capillary architecture. For example, in the hepatic lobule, which is the fundamental structural unit of the liver, sinusoids radiate outward from the central vein (26, 27). In contrast, capillaries in the kidney glomerulus are densely packed within a spherical structure (28, 29). Thus, the vascular geometry and 3D organization of each organ are specifically adapted to its function (30). Reproducing such microvascular networks is essential for supporting proper cell metabolism.

Bioprinting technologies based on stereolithography or injection molding have been utilized for the *in vitro* construction of tissues with predefined shapes (31, 32). However, these technologies are typically used to fabricate relatively large tissues exceeding 100 µm in diameter, and their limited spatial resolution prevents their application in capillary and microvessel construction. Recent advances in multiphoton lithography-based bioprinting have enabled highly flexible, high-resolution 3D fabrication (33). This laser-based method allows for the creation of microchannels with arbitrary 3D geometries and micrometer-level precision. Using this technology, Enrico et al. (34) fabricated hydrogel channels with diameters ranging from 20 to 60 µm and seeded them with vascular endothelial cells, which led to immediate lumen formation along the channels. Arakawa et al. (35) created hydrogel channels with diameters ranging from 7 to 40 µm, incorporating a central neck structure, and successfully induced vessel formation along the channels while also replicating the pathological behavior of malaria nematode-infected erythrocytes. Similarly, Rayner et al. (36) used this approach to replicate organ-specific vascular structures. Recently, Salvadori et al. (37) created straight microvascular structures with diameters ranging from 50 to 110 µm and demonstrated their physiological and inflammatory relevance.

Most of the vessels generated in these studies resemble small arterioles, with diameters of several tens of micrometers, and do not replicate capillaries below 20 µm, which are responsible for substance exchange *in vivo*. Furthermore, endothelial cells were not directly seeded into capillary-scale microchannels; instead, cells were initially introduced into larger-diameter channels, followed by laser ablation to damage the endothelial monolayer and induce capillary-scale angiogenesis (35, 36). This approach imposes restrictions on the timing and placement of cell seeding. Moreover, the mechanisms underlying efficient vascular formation using capillary-scale microchannel templates remain poorly understood. Therefore, to achieve more efficient *in vitro* capillary construction with predefined geometries, it is essential to elucidate the process of vascular formation in capillary-scale microchannels.

In this study, we investigated the process of microvessel formation along hydrogel microchannel templates. Although synthetic materials such as polyethylene glycol diacrylate and gelatin methacryloyl are commonly used in bioprinting (38–41), we selected natural hydrogel materials, such as collagen and fibrin, due to their biochemical and mechanical properties, which more closely resemble those of native tissues. Microchannels were fabricated within the hydrogels using femtosecond laser ablation via multiphoton absorption. First, we optimized the parameters of multiphoton lithography required to fabricate hollow channels within hydrogels, which serve as templates for capillary and microvessel construction. Next, we developed a method for constructing vascular structures in predefined geometries. We observed immediate vascular formation along 50 μm channels. However, under static conditions, endothelial cells failed to form microvessels along 20 μm channels. These findings suggests that as channel diameter decreases to a scale comparable to that of individual cells, the mode of vascular formation shifts significantly. Our results demonstrated that flow stimulation significantly enhanced vascular formation, whereas vascular formation failed under static conditions. Vessel formation was assessed by tracking individual cells using time-lapse microscopy. Furthermore, immunofluorescence staining revealed differences in the expression of proteins associated with cell–cell adhesion and migratory ability. Notably, our findings suggest that under flow conditions, vascular formation occurs through collective cell behavior with enhanced cell–cell adhesion, whereas endothelial-to-mesenchymal transition (EndMT) may occur under static conditions. These findings represent a significant step forward in the *in vitro* construction of organ-specific capillary networks and the development of more complex vascularized architectures with predefined shapes.

## Materials and Methods

### Preparation of microfluidic devices

The fabrication of the microfluidic devices used in this study was previously described (42). Briefly, polydimethylsiloxane (PDMS; Silgard 184, Dow Corning, Midland, MI, USA) was fabricated by soft lithography and plasma-bonded with a coverglass. Then, microchannels were coated with 1 mg/ml poly-D-lysine solution (Sigma-Aldrich, St. Louis, MO, USA) and rinsed twice with sterile deionized water. The gel channels were designed with widths of 1,000 and 600 μm, respectively, and a height of 150 μm (Fig. 1A).

**Fig. 1.**
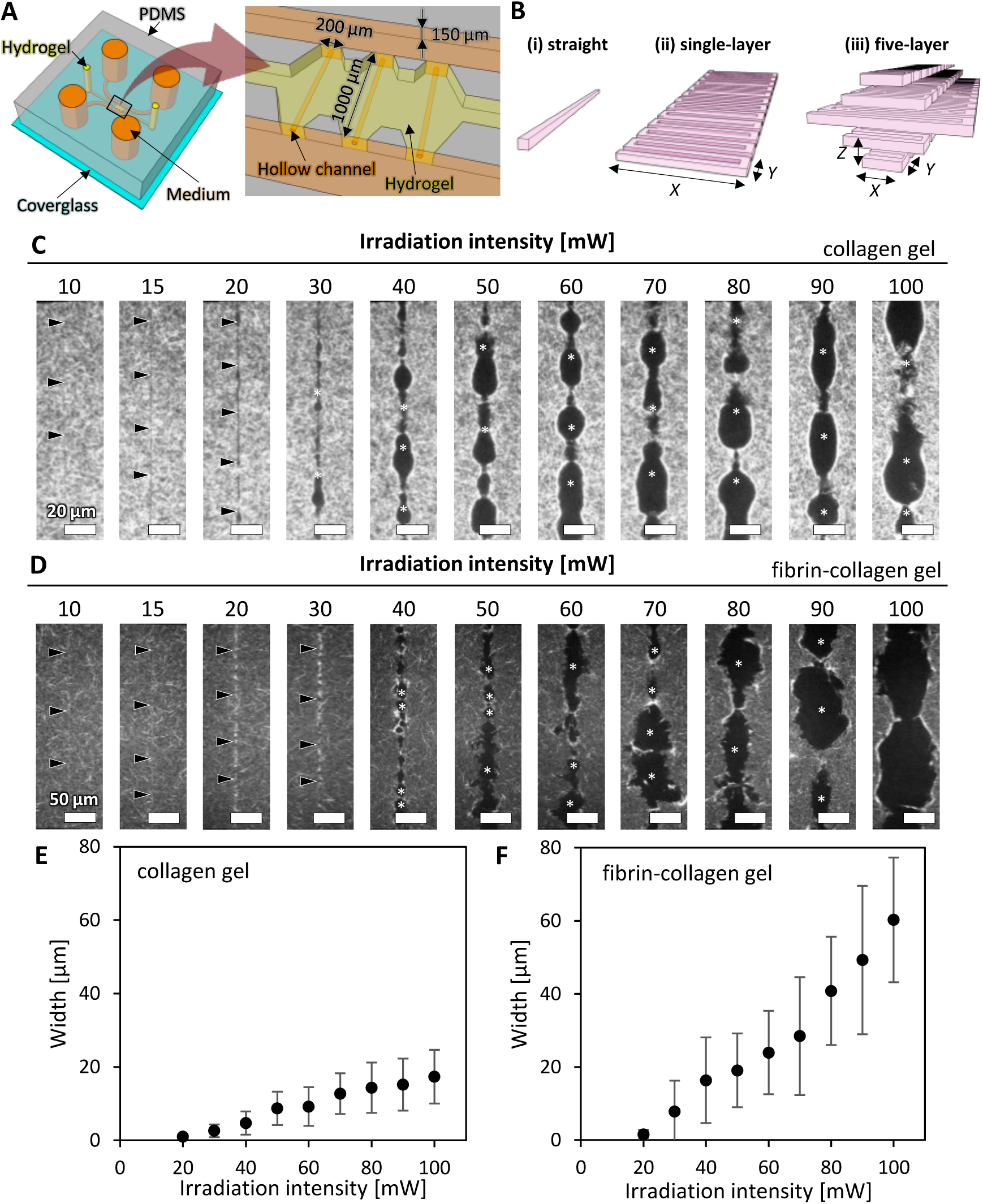
Schematic illustrations of a microfluidic device and fabrication of hollow structures along the straight trajectory at various irradiation intensities. (A) A microfluidic device consisting of three parallel microchannels: a central channel for hydrogel and two adjacent channels for culture medium. Hollow channels were fabricated within the hydrogel. (B) Laser irradiation trajectories: (i) straight, (ii) single-layer zigzag, and (iii) five-layer zigzag patterns. Laser irradiation parameters: *X* — reciprocating width, *Y* — reciprocating interval, *Z* — layer interval. (C) Confocal reflection images of hollow structures in collagen gel. Scale bars, 20 μm. Arrowheads indicate the structures fabricated by photoablation. Asterisks indicate the structures fabricated by cavitation molding. (D) Confocal reflection images of hollow structures in fibrin-collagen gel. Scale bars, 50 μm. (E) Quantification of the width of fabricated hollow structures in collagen gel. (F) Quantification of the width of fabricated hollow structures in fibrin-collagen gel. Data are shown as the mean ± SD. n=51.

### Fabrication of hollow channels within hydrogels by multiphoton ablation

Hollow channels were fabricated inside the microfluidic devices placed on an XYZ motorized stage, using a femtosecond laser source (Carbide, Light Conversion, Lithuania) coupled with an optical parametric amplifier (Orpheus, Light Conversion, Lithuania). The fabrication geometry was initially designed in STL format using 3D CAD software. The data were then converted into trajectory files for stage control. In this study, straight, single-layer zigzag, and five-layer zigzag scanning patterns were selected based on the intended 3D structures, as illustrated in Fig. 1B. Ultrashort pulse laser beams (wavelength: 700 nm, pulse width: approximately 260 fs) from the femtosecond laser system were focused into the hydrogel within the microfluidic devices using an objective lens (M Plan Apo NIR HR 100×, Mitsutoyo, Japan). To ensure precise fabrication of hollow channels, a two-camera system, consisting of a high-magnification microscope and a large-field-of-view microscope, was used to simultaneously monitor both the laser irradiation area and the overall processed device. Fabrication was performed by scanning the laser with a 1-μm step size using an XYZ motorized stage, following the precomputed trajectory data files. The fabricated structures were subsequently examined using a confocal microscope.

### Cell culture

Human umbilical vein endothelial cells (HUVECs; Lonza) were obtained commercially and cultured in an endothelial growth medium (EGM-2; Lonza). The cells were maintained in a humidified incubator at 37°C with 5% CO_2_ and used for experiments at passage 4.

### Construction of template-guided microvessels and perfusion culture in a microfluidic device

First, a rat-tail type-I collagen solution (8 mg/ml, pH 7.4; BD Biosciences, San Jose, CA, USA) was injected into a gel port (Fig. 1A). The samples were placed in a humidified incubator at 37°C with 5% CO_2_ for 30 min to allow gelation. Subsequently, HUVECs were seeded onto one side of the gel surface by injecting a 20 µl cell suspension (5.0 × 10^6^ cells/ml) into the medium port (Fig. 4A, B). Immediately after seeding, the device was tilted to allow the cells to settle onto the gel surface by gravity. After seeding, the samples were returned to the incubator for 2 h to promote cell adhesion.

Reservoirs were connected to the outlets of the microchannels to establish flow. The cells were cultured under static or flow conditions. In static conditions, an equal volume of medium was placed in the reservoirs. In flow conditions, a gravity-driven flow was generated by maintaining a hydrostatic pressure difference of 5 or 10 mmH_2_O (Fig. 4B). The culture medium was replaced daily to restore the flow. Although the hydrostatic pressure difference gradually decreased over time, it was confirmed that the pressure difference did not equilibrate within 24 h between medium changes (6).

### Time-lapse microscopy

For the time-lapse experiment, images were captured continuously over 24 h from day 2 to day 3 after cell seeding into the devices. Phase-contrast images were acquired every 15 min using a 20× objective lens on a time-lapse microscope (ECLIPSE Ti-E; Nikon, Japan) equipped with a stage top incubator (Tokai Hit, Japan). The imaging positions were pre-registered, ensuring that all blood vessels within the device were automatically imaged. Immediately following the final imaging session, the cells were fixed with 4% paraformaldehyde.

### Immunofluorescence staining of cultured cells

The cells were fixed with 4% paraformaldehyde for 15 min at room temperature. Following fixation, the cells were rinsed twice with phosphate-buffered saline (PBS) and treated with 0.1% Triton X-100 and BlockAce (Dainippon Pharmaceutical, Osaka, Japan) for 1 h to permeabilize the membranes and block nonspecific staining. After two additional PBS rinses, the cells were incubated overnight at 4°C with primary antibodies: sheep anti-PECAM-1 (1:100 dilution; AF806, R&D Systems) and mouse anti-vimentin (1:200 dilution, SC6260; Santa Cruz). Subsequently, the samples were incubated overnight with secondary antibodies: Alexa Fluor 555-conjugated anti-mouse IgG (1:200 dilution; Invitrogen) and Alexa Fluor 488-conjugated anti-sheep IgG (1:200 dilution; Invitrogen). Z-stack fluorescence images were acquired using a confocal laser-scanning microscope (LSM700; Carl Zeiss, Hallbergmoss, Germany). Projection and cross-sectional images were generated using ImageJ (National Institutes of Health, Bethesda, MD).

### Velocity field analysis

The velocity field of collective cellular migration within the collagen channel was analyzed using a conventional particle imaging velocimetry (PIV) algorithm, implemented in the PIVlab software (36). Sequential time-lapse images were processed with a 32 × 32-pixel interrogation area and a 16-pixel step size to detect local movements. The resulting velocity field along the collagen channel was visualized in HSV color space, where the hue represented the orientation and the saturation indicated the magnitude of local movement within the exploratory window. The value component was fixed at 1. The generated colored mosaic images representing the velocity field at each time point were overlaid onto the original time-lapse image, as shown in Supplementary Movies 3 and 4.

### Statistical analyses

All experiments were repeated at least two to three times to ensure the reproducibility of the results. Data are presented as the means ± SD.

## Results

### Fabrication of hollow channels within hydrogels by multiphoton lithography

We investigated the parameters of multiphoton lithography required to fabricate microscale hollow channels within hydrogels, which serve as templates for constructing capillaries and microvessels. Two types of hydrogels were used: collagen gel and fibrin-collagen gel, both of which are widely utilized as scaffolds for *in vitro* microvascular formation.

First, we investigated the effect of laser irradiation intensity using a straight-line irradiation trajectory (Fig. 1B(i)). In collagen gels, laser irradiation intensities below 20 mW produced narrow cavities along the irradiation path (Fig. 1C, arrowheads), likely due to photoablation. While processing marks were observed at 10 and 15 mW (Fig. 1C, arrowheads), continuous removal of the collagen gel was not achieved. In contrast, an intensity of 20 mW resulted in the continuous removal of the collagen gel (Fig. 1C, arrowheads).

On the other hand, relatively large cavities began to form at an intensity of 30 mW, likely due to cavitation molding (Fig. 1C, asterisks). As the irradiation intensity increased, the fabrication mechanism transitioned from photoablation to cavitation molding (Fig. 1C), consistent with a previous study (34). Increasing irradiation intensities resulted in the formation of larger structures within collagen gels, with an intensity of 100 mW producing cavities approximately 20 µm in width (Fig. 1E).

Similarly, in fibrin-collagen gels, laser irradiation intensities below 30 mW produced narrow cavities along the irradiation path (Fig. 1D, arrowheads), likely due to photoablation. Intensities of 20 and 30 mW led to the continuous removal of the fibrin-collagen gel (Fig. 1D, arrowheads). In contrast, intensities above 40 mW produced relatively large cavities (Fig. 1D, asterisks). The surfaces of these cavities were rougher compared to those in collagen gels (Fig. 1D). Increasing irradiation intensities produced larger structures within fibrin-collagen gels, with an intensity of 100 mW creating cavities approximately 60 µm in width (Fig. 1F). These results indicate that both collagen and fibrin-collagen gels can be precisely fabricated along the irradiation path at intensities of <20 mW.

Next, we further investigated laser fabrication conditions to construct cylindrical hollow channels with a diameter of 20 µm, which is comparable to the scale of capillaries and microvessels. We extended the straight-line, one-dimensional irradiation trajectory (Fig. 1B(i)) to a planar, 2D trajectory (Fig. 1B(ii)), and then layered it (Fig. 1B(iii)) to establish conditions for constructing 3D structures, such as hollow channels.

Each layer was fabricated using zigzag irradiation trajectories, with the reciprocating interval (*Y*), laser irradiation frequency, and stage speed adjusted to ensure stable fabrication along the trajectories (Supplementary Figs. 1, 2, 3). Based on these results, we selected a reciprocating interval of 2 µm. We then examined the irradiation intensities using single-layer and five-layer trajectories to fabricate collagen gels and fibrin-collagen gels (Fig. 2A). In the single-layer trajectory, unprocessed hydrogel remained at an intensity of 10 mW in both types of hydrogels (Fig. 2B, C). An intensity of 15 mW allowed for successful fabrication of the hydrogel along the irradiation trajectory. However, at 20 mW, fabrication resulted in an unstable width, likely due to the generation of cavitation bubbles.

**Fig. 2.**
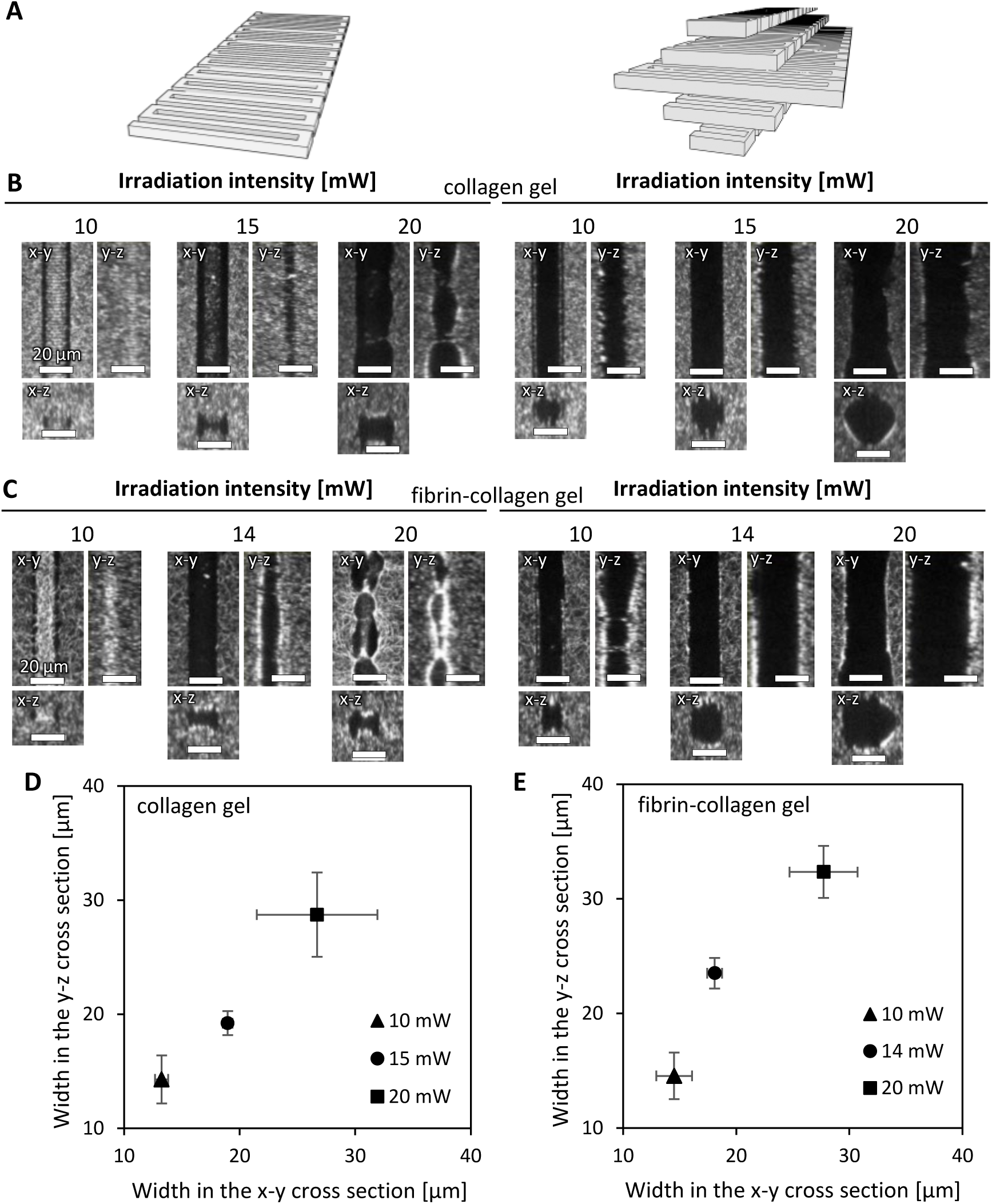
Hollow structures fabricated along single-layer and five-layer trajectories at various irradiation intensities. (A) Single-layer and five-layer laser irradiation trajectories. (B) Confocal reflection images of the hollow structures in collagen gel fabricated along the single-layer or five-layer trajectories. Scale bars, 20 μm. (C) Confocal reflection images of the hollow structures in fibrin-collagen gel fabricated along the single-layer or five-layer trajectories. Scale bars, 20 μm. (D) Quantification of the width of the hollow structures in collagen gel fabricated along the five-layer trajectory. (E) Quantification of the width of the hollow structures in fibrin-collagen gel fabricated along the five-layer trajectory. Data are shown as the mean ± SD. n=51.

For the five-layer trajectory, we investigated the appropriate layer interval (*Z*) and selected 2 µm to fabricate hollow channels (Supplementary Fig. 4). At this layer interval, an intensity of 10 mW produced slightly smaller hollow channels than those with a diameter of 20 µm in collagen gels (Fig. 2B), with a rougher luminal surface. In contrast, an intensity of 15 mW generated hollow channels with a diameter of approximately 20 µm, closely matching the width of the irradiation trajectory, and a smoother luminal surface. An intensity of 20 mW resulted in hollow channels larger than the width of the irradiation trajectory (Fig. 2B), likely due to the generation of cavitation bubbles. Similarly, in fibrin-collagen gels, an irradiation intensity of 14 mW produced hollow channels with a dimeter of approximately 20 µm (Fig. 2C).

Quantitative analysis of the fabricated widths in the x-y and y-z cross-sections was performed for the five-layer trajectory. In collagen gels, the fabricated widths in the x-y and y-z cross-sections were 18.7±0.6 µm and 18.7±1.4 µm, respectively, at an intensity of 15 mW (Fig. 2D), while in fibrin-collagen gels, the widths were 18.2±0.7 µm and 23.2±1.8 µm at an intensity of 14 mW (Fig. 2E). Based on these results, the irradiation intensities that produced hollow channels with a diameter of 20 µm were determined to be 15 mW for collagen gels and 14 mW for fibrin-collagen gels. Hollow channels with a diameter of 50 µm were also fabricated using these conditions with some modifications.

### Vascular formation guided by channels with a diameter of 50 μm

Many studies have reported methods for forming luminal structures within hydrogels and creating blood vessels using these templates, particularly focusing on diameters in the range of several hundred micrometers (43). Although this study aims to construct blood vessels at the scale of capillaries and microvessels, we first constructed blood vessels using a template with a diameter of 50 μm in collagen gels as a control experiment (Fig. 3). When the suspension of HUVECs was introduced into the microfluidic channel, the fabricated hollow channel was rapidly filled with the cells (Fig. 3A, after). The cells then attached to the luminal surface of the hollow channel within 2 h, with some cells spreading (Fig. 3A, 2 h). By day 1, almost the entire channel was covered with cells (Fig. 3A, day 1), and the vascular structures remained intact on day 2 (Fig. 3A, day 2). Confocal fluorescence images of the cells on day 2 showed that they covered the entire luminal surface of the hollow channel, resulting in the formation of vascular tissue with a continuous lumen (Fig. 3B). These vascular structures were constructed in static culture, where cell adhesion and spreading occurred similarly to that in 2D culture. These results indicate that we successfully constructed blood vessels guided by channels with a diameter of 50 μm within two days using our experimental system.

**Fig. 3.**
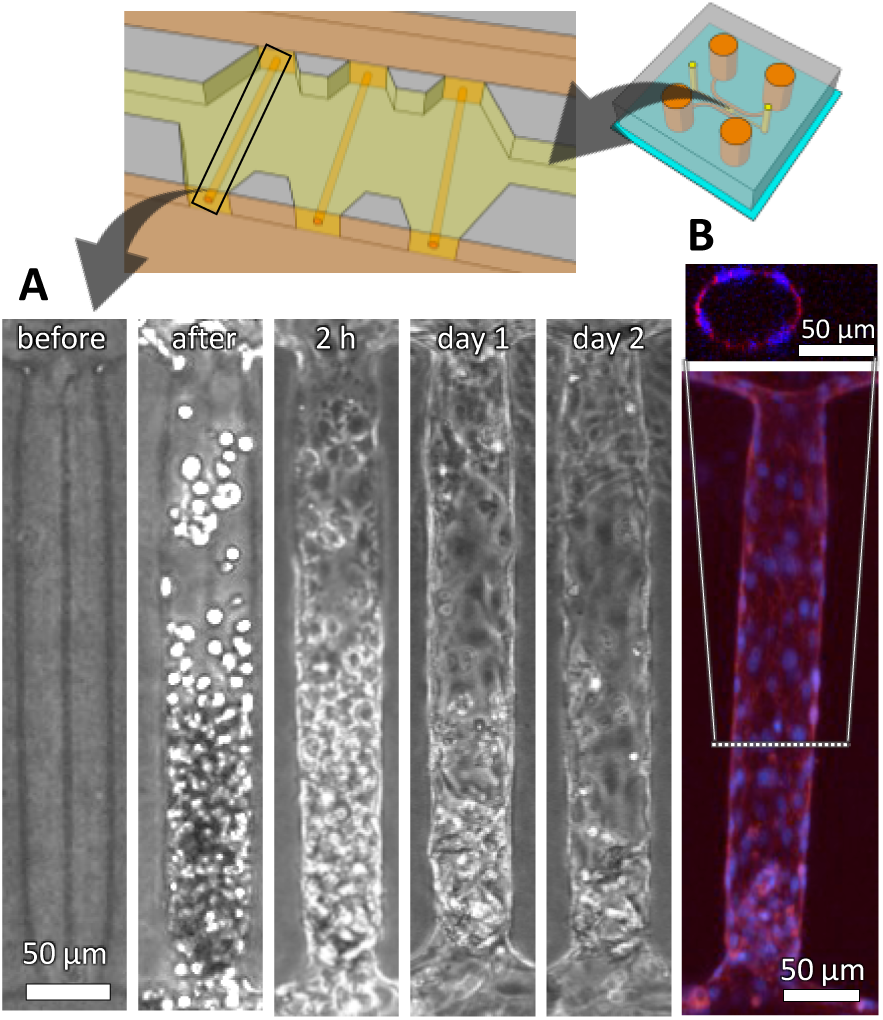
Vascular formation guided by a hollow structure with a 50 μm diameter under static conditions. (A) Representative phase-contrast images of HUVECs during vascular formation. Scale bars, 50 μm. (B) A fluorescent projection image of a straight blood vessel generated from z-stack confocal images and a corresponding cross-sectional image. HUVECs were fixed on day 2 and stained for F-actin (phalloidin) and nuclei (Hoechst 33342). Scale bars, 50 μm.

### Vascular formation guided by channels with a diameter of 20 μm

We then constructed blood vessels using a template with a diameter of 20 μm, which is similar in scale to capillaries and microvessels, in collagen gels. First, cells were cultured under static conditions (Fig. 4A). Upon seeding the cells, some cells entered the hollow channel, while others became clogged at the entrance (Fig. 4C, day 0, arrowheads). By day 1, more cells migrated into the channels, but they were neither extending nor covering the luminal surface (Fig. 4C, day 1). From day 3 to day 5, some cells began to extend and cover the luminal surface; however, a continuous tissue was not formed. There were areas indicating that the tissue was narrowing locally and becoming discontinuous (Fig. 4C, days 3, 5, asterisks). On day 7, the number of the cells remaining within the channels decreased (Fig. 4C, day 7). A corresponding fluorescence image of the cells on day 7 showed that they were distributed only in small regions within the hollow channel and were not spreading on the luminal surface (Fig. 4C, day 7). A cross-sectional projection image revealed that a single cell adhered to multiple surfaces of the channel under static conditions (Fig. 4E, static). Although blood vessels were successfully constructed using the template with a diameter of 50 μm, we failed to construct blood vessels using the template with a diameter of 20 μm under static conditions.

**Fig. 4.**
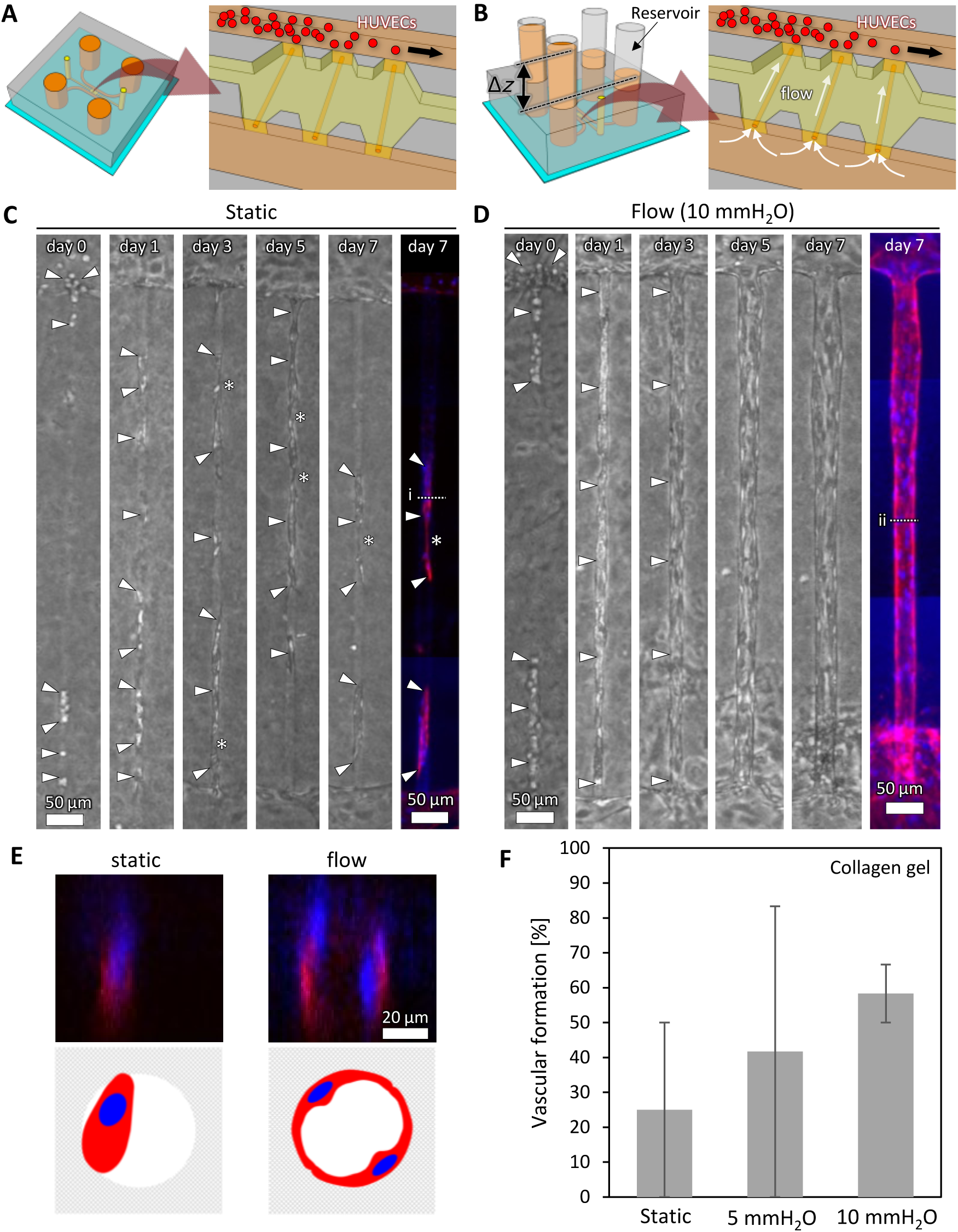
Vascular formation guided by a hollow structure with a 20 μm diameter. (A) Schematic illustrations of a microfluidic device under static conditions. (B) A microfluidic device connected to tube reservoirs. A liquid level difference (Δ*z*) generated flow through the hydrogel from a microchannel without HUVECs to one with HUVECs. White arrows indicate the flow direction. (C) Representative phase-contrast images of the cells from day 0 to day 7 under static conditions, and a corresponding fluorescence image stained for F-actin (phalloidin) nuclei (Hoechst 33342) on day 7. A projection image was generated from z-stack confocal images. Arrowheads indicate the cells introduced into the hollow structure. Asterisks indicate where multicellular structures are thinning. Scale bars, 50 μm. (D) Representative phase-contrast images under flow conditions, and a corresponding fluorescence image stained for F-actin (phalloidin) nuclei (Hoechst 33342) on day 7. Scale bars, 50 μm. (E) Cross-sectional projection images generated from z-stack confocal images corresponding to lines i and ii in C and D, along with schematic illustrations of cells within hollow channels. (F) Vascular formation rate in collagen gel. Hollow structures with >50% cellular occupancy on day 7 were considered as vascular formation. Data are shown as the mean ± SD. N=2. n=6.

To promote vascular formation along channels with a diameter of 20 μm, interstitial flow, known to enhance angiogenesis (17), was applied in the direction opposite to vessel elongation. A pressure difference of 10 mmH_2_O across the hydrogel scaffold generated flow through the channel (Fig. 4B). By day 1, many cells migrated into the channels, with some cells spreading and covering the luminal surface (Fig. 4D, day 1). By day 7, the entire luminal surface was covered with cells, leading to the formation of vascular structures guided by the channels (Fig. 4D, days 3, 5, 7). Additionally, the constructed vessels thickened from days 3 to 7. A cross-sectional projection image generated from z-stack confocal images revealed that cells spread along the luminal surface, forming a continuous lumen under 10 mmH_2_O flow conditions (Fig. 4E, flow). In the FITC-dextran perfusion experiment, the continuous lumen which covered by endothelial cells presented barrier function (Supplementary Fig. 5). When comparing the rates of vascular formation under static conditions, 5 mmH_2_O, and 10 mmH_2_O flow conditions, the highest rate was observed under 10 mmH_2_O flow conditions (Fig. 4F). Representative images of cells cultured for 7 days under each condition, in both collagen and fibrin-collagen gels, are provided in supplementary materials (Supplementary Figs. 6, 7). These results demonstrate that vascular formation along channels with a diameter of 20 μm was successfully induced under flow conditions.

### Evaluation of the vascular formation process in static and flow conditions

To further investigate the vascular formation process along channels with a diameter of 20 μm, we performed phase-contrast time-lapse imaging under both flow and static conditions. Under static conditions, cells initially located around the entrance and inside the hollow channel connected to form a continuous structure (Fig. 5A, 0 h). However, over time, this structure locally thinned and eventually split into two groups due to disrupted cell–cell adhesion (Fig. 5A, 6 h, asterisks). Consequently, the leading group of cells became isolated within the channel (Fig. 5A, 12 h), and further fragmentation into smaller cell clusters was observed (Fig. 5A, 24 h). Tracking individual cells in the time-lapse movie (Supplementary Movie 1) also revealed that some cells migrated out of the channel (Fig. 5C, arrows), further highlighting the instability of vascular formation under static conditions.

**Fig. 5.**
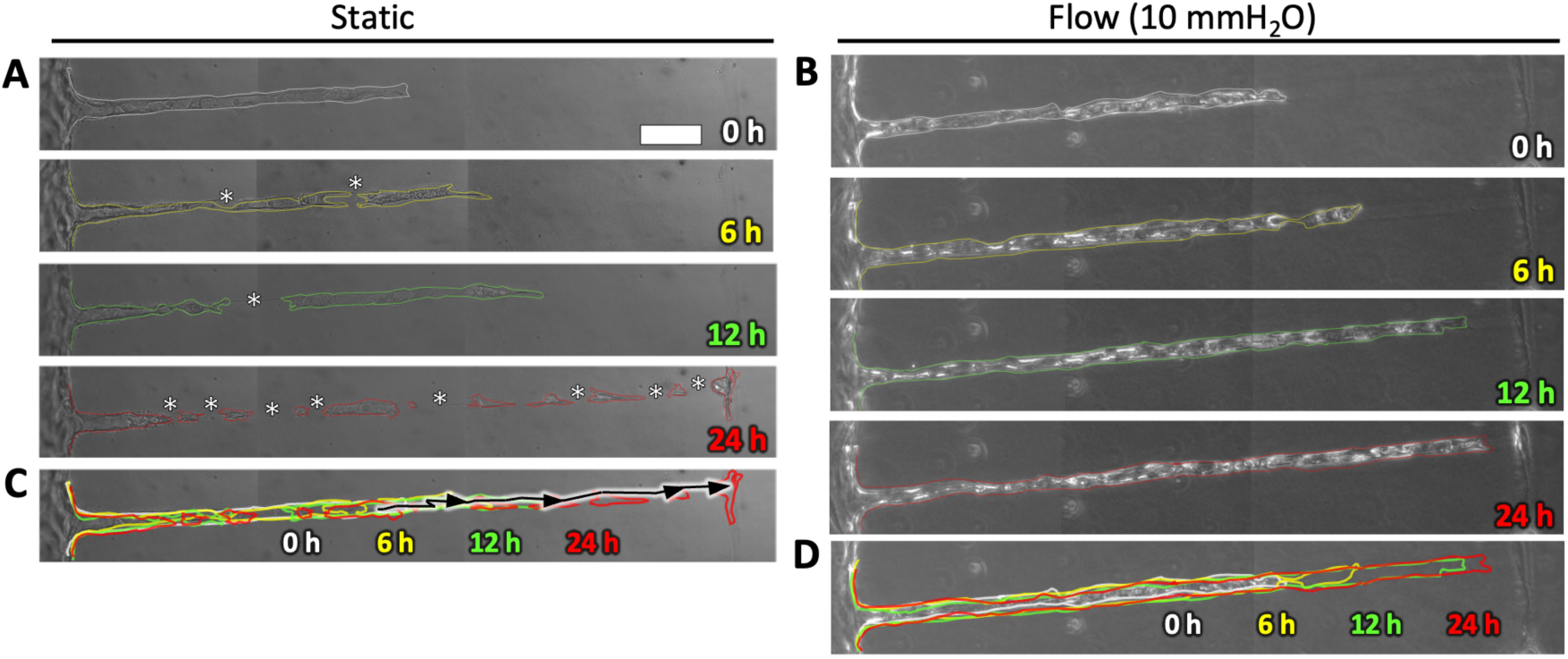
Vascular formation process under static and flow conditions. (A, B) Time-lapse phase-contrast images were captured from days 2 to 3 to investigate the behavior of HUVECs seeded in a hollow structure with a 20 μm diameter. Representative phase-contrast images show the cells at 0 h, 6 h,12 h, and 24 h under static and 10 mmH_2_O flow conditions. (C, D) Superimposed image of cellular outlines under static and flow conditions. The black line indicates the trajectory of a single cell migrating out of the hollow structure. Scale bar, 50 µm.

In contrast, under flow conditions, cells initially located around the entrance and inside the hollow channel collectively migrated and gradually covered the luminal surface of the channel (Fig. 5B, 0, 6, 12 h). This structure continued to elongate along the channel without disruptions of cell–cell adhesion and eventually covered most of the channel (Fig. 5B, 24 h), leading to the formation of a continuous vascular structure (Fig. 5D). Time-lapse movie under flow conditions is provided in the supplementary materials (Supplementary Movie 2). Furthermore, image correlation analysis of migration direction confirmed the directional bias of cell movement, consistent with observations from the time-lapse movies under both static and flow conditions (Supplementary Movies 3, 4). These findings suggest that cell–cell adhesion and cell motility play crucial roles in vascular formation under different environmental conditions.

**Fig. 6.**
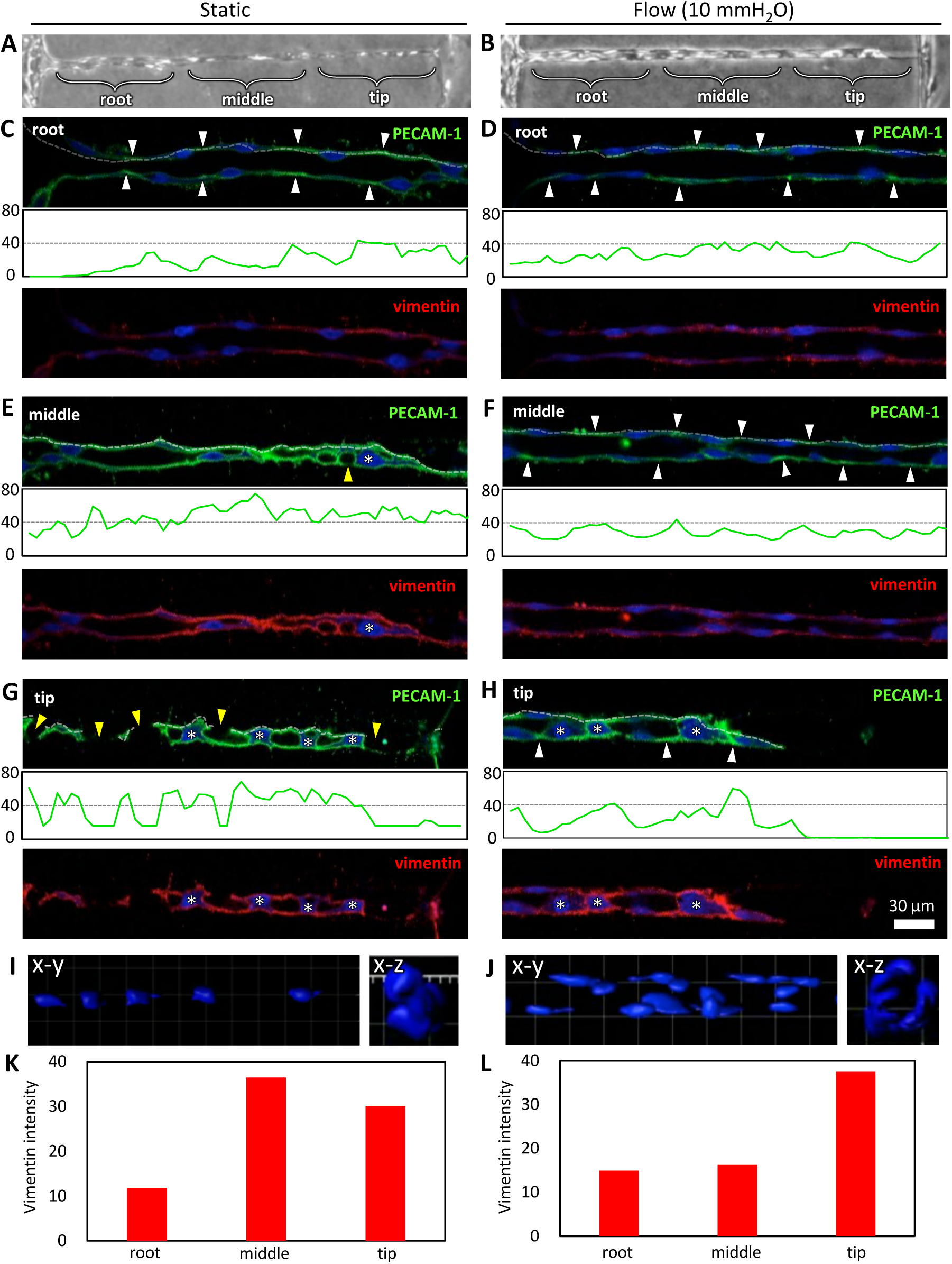
Evaluation of protein expression during vascular formation in a 20 μm diameter channel. (A, B) Phase-contrast images of HUVECs on day 3 under static and flow conditions. Vascular structures were categorized into root, middle, and tip regions. (C–H) Cross-sectional immunofluorescence images of cells within straight microchannels in the root, middle, and tip regions. Cells were fixed on day 3 and stained for PECAM-1 (green), vimentin (red), and nuclei (Hoechst 33342, blue). White arrowheads indicate PECAM-1 localized at cell–cell borders, while yellow arrowheads mark disconnected cell–cell borders. Asterisks indicate nuclei occupying the luminal space, misaligned with the luminal surface of the hollow structure. Scale bar, 30 μm. Fluorescence intensity distribution along the dotted line in PECAM-1 images. (I, J) 3D reconstruction images of cell nuclei in the tip and root regions under static and flow conditions. (K, L) Quantification of vimentin intensity in the root, middle, and tip regions under static and flow conditions.

Since reduced cell–cell adhesion and enhanced cell motility are the characteristic features of EndMT, we performed immunofluorescence staining to examine proteins involved in this process (44, 45). Specifically, we analyzed the expression of PECAM-1 as a marker of cell–cell adhesion and vimentin as an indicator of EndMT (46). The vascular structures formed along the channel were categorized into three distinct regions: the “root region,” where the vascular structure connected to a microfluidic channel containing endothelial cells; the “middle region,” located at the center of the channel; and the “tip region,” representing the elongating front of the vascular structure (Fig. 6A, B).

In the root region, cells adhered to and spread along the luminal surface of the hollow channel under both flow and static conditions, with their nuclei aligned along the luminal surface, leading to the formation of a vascular lumen (Fig. 6C, D). PECAM-1 expression was localized between cells, likely at cell–cell junctions (Fig. 6C, D, white arrowheads).

In contrast, in the middle region, PECAM-1 did not localize at cell–cell junctions and instead highly expressed across the entire cell membrane under static conditions (Fig. 6E). Additionally, the nuclei were misaligned with the luminal surface, occupying most of the hollow channel, and no vascular lumen was formed (Fig. 6E, asterisks). Moreover, disrupted cell–cell junctions were observed in some areas (Fig. 6E, yellow arrowheads). Under flow conditions, however, the nuclei aligned along the luminal surface, and a vascular lumen was formed (Fig. 6F). As in the root region, PECAM-1 localized at cell–cell junctions (Fig. 6F, white arrowheads).

In the tip region, the nuclei were misaligned with the luminal surface, occupying most of the hollow channel, and no vascular lumen was formed under static conditions, similar to the middle region (Fig. 6G, asterisks). Additionally, many disrupted cell–cell junctions were observed (Fig. 6G, yellow arrowheads). In contrast, under flow conditions, PECAM-1 was localized between cells, with cells that spread along the luminal surface showing aligned nuclei, while cells that did not spread exhibited misaligned nuclei (Fig. 6H).

Fluorescent 3D reconstruction images of cell nuclei clearly demonstrate that, under static conditions, the nuclei misaligned along the luminal surface and occupied the luminal space of the channel in the tip region (Fig. 7I). In contrast, under flow conditions, the nuclei were well-aligned along the luminal surface of the channel, forming continuous lumens in the root region (Fig. 7J).

**Fig. 7.**
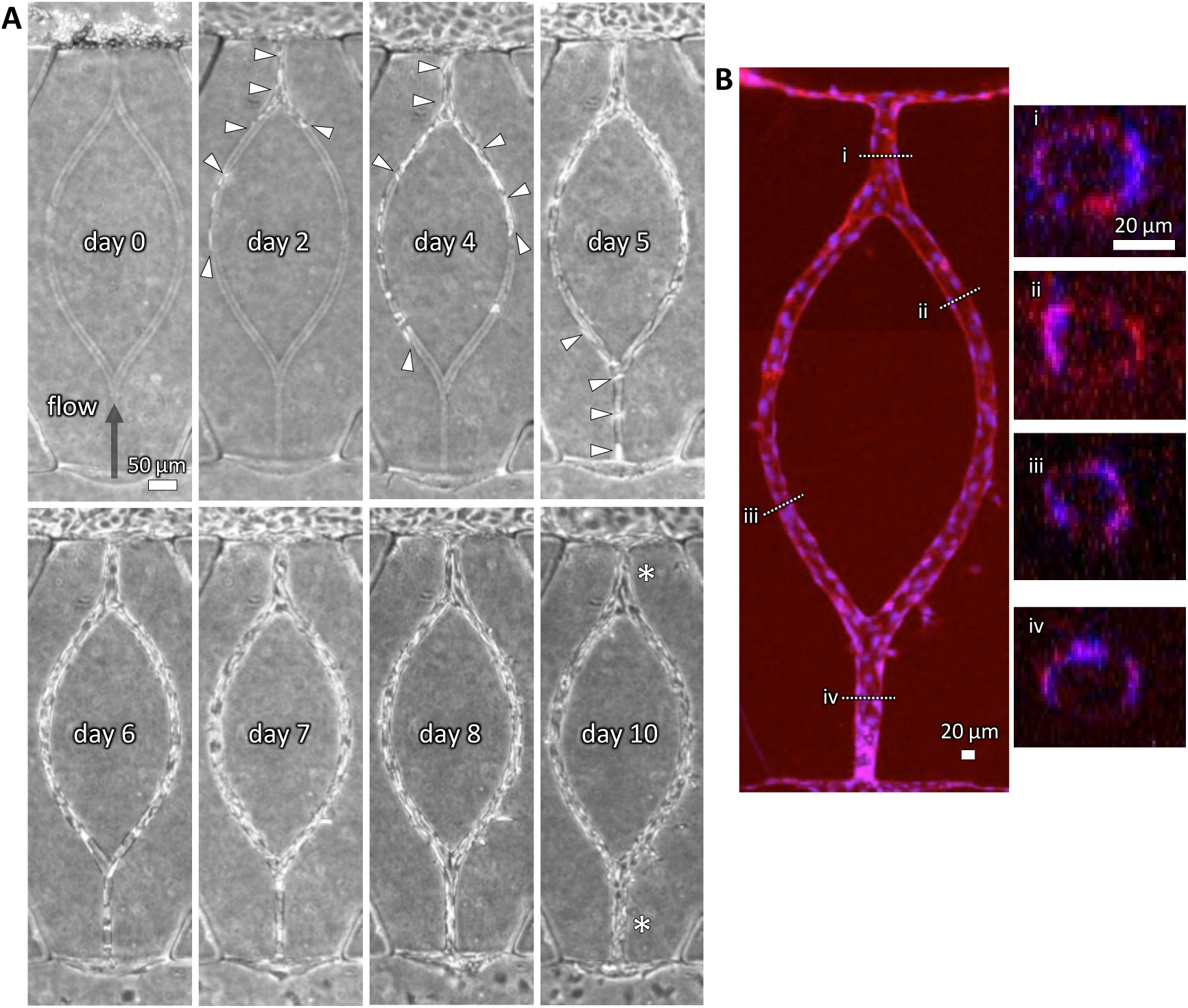
Vascular formation guided by a bifurcated hollow structure under flow condition. (A) Corresponding phase-contrast images of vascular formation in collagen gel from days 0 to 10. Arrowheads indicate the cells within the bifurcated hollow structure. The cells were cultured under flow conditions, with the flow direction indicated by the black arrow. (B) A corresponding fluorescence image on day 10 showing F-actin (red, phalloidin) and nuclei (blue, Hoechst 33342). The projection image was generated from z-stack confocal images. Cross-sectional images corresponding to lines i–iv are also shown. Scale bars: 50 µm (A), 20 µm (B), 20 µm (i).

The expression of vimentin was weak in cells that spread along the luminal surface and formed a vascular lumen (Fig. 6C, D, F), while strong expression was observed in cells that failed to form a vascular lumen and had nuclei misaligned with the luminal surface (Fig. 6E, G, H). Quantification of vimentin intensity at the root, middle, and tip regions confirmed these findings (Fig. 6K, L).

### Vascular formation guided by a bifurcated hollow channel under flow conditions

Multiphoton lithography offers the capability to fabricate hollow channels with predefined shapes and sizes by precisely controlling the trajectory of laser irradiation, enabling the construction of microvessels in any desired geometries. As a proof of concept, we fabricated a branched-loop channel with a diameter of 20 μm within a collagen gel and cultured HUVECs to construct a bifurcated microvessel. To enhance vascular formation along the channel, cells were cultured under flow conditions, with the flow direction set opposite to that of vascular elongation (Fig. 7A, arrow).

Phase-contrast images revealed that cells migrated into the channel, with some cells spreading and covering the luminal surface around the entrance from days 0 to 5 (Fig. 7A, days 0, 2, 4, 5). As the cells gradually migrated along the channel, they eventually covered the entire luminal surface by day 8, leading to the formation of bifurcated microvessels (Fig. 7A, day 8). Furthermore, from days 8 to 10, the vessel diameter increased at the junction where the two branches converged (Fig. 7A, day 10, asterisks).

Confocal fluorescence images on day 10 confirmed that the vascular tissue had a continuous lumen (Fig. 7B). In the branched region, the lumen diameter was approximately 20 μm, while the lumens before and after the convergence measured approximately 30 μm each (Fig. 7B).

## Discussion

### Two distinct vascular formation processes depending on the vessel diameter

In this study, we successfully fabricated hollow channels within a hydrogel using multiphoton lithography. Channels with a diameter of 20 μm were designed as templates for constructing capillary-scale microvessels, while those with a diameter of 50 μm were created to construct larger vessels. Notably, we observed significant differences in vascular formation processes between these two templates. For channels with a diameter of 50 μm, vessels were rapidly formed under static conditions within 2 days. This formation process is consistent with previous studies on vascular formation using collagen microchannels created by placing a microneedle during gelation (43). The size of these microchannels is determined by the diameter of microneedles, typically ranging from 200 to 500 μm. In contrast, the multiphoton lithography approach used in this study enabled the fabrication of collagen microchannels with a diameter of at least 20 μm, a size that cannot be achieved with the microneedle-based method.

In contrast to channels with a diameter of 50 μm, endothelial cells failed to form microvessels guided by channels with a diameter of 20 μm under static conditions. This result suggests that the mode of vascular formation shifts as the diameter of the channel decreases to a scale comparable to that of individual cells. As shown in Fig. 4, many cells accumulated around the entrance of the channel rather than entering it, with only a small number successfully introduced into the channels. To form microvessels, the cells needed to migrate into the channel and cover the luminal surface. Our previous study demonstrated that interstitial flow applications significantly promote the extension of angiogenic sprouts against the flow direction (17). Based on this finding, we applied flow through the channel to facilitate cell migration, which successfully promoted cell entry and resulted in the formation of microvessels along the channel.

We demonstrated that flow application is essential for the successful vascular formation along microchannels with a diameter of 20 μm, which falls within the range of capillaries and microvessels. This finding is significant because microvessel construction along microchannels is a critical step in various bioprinting approaches and the recellularization of decellularized tissues and organs. For instance, constructing capillary-scale microvessels, such as sinusoids, in decellularized livers remains a significant challenge (47).

### The mechanism of successful vascular formation in 20 µm diameter channels under flow conditions

Vascular formation within 20 µm diameter channels was significantly enhanced under flow conditions compared to static conditions (Figs. 4 and 5). Time-lapse imaging revealed that flow conditions facilitated collective migration along the channels, leading to the elongation of nascent vascular tissues (Figs. 5B). In contrast, static conditions were characterized by prominent individual cell migration and disrupted cell–cell junctions, which resulted in the fragmentation of cell clusters (Figs. 5A, asterisks). These observations suggest that EndMT may occur under static conditions. EndMT is a complex biological process in which endothelial cells adopt a mesenchymal phenotype displaying typical mesenchymal cell morphology and functions, including acquisition of cellular motility and contractile properties. Endothelial cells undergoing EndMT lose the expression of endothelial cell-specific proteins (44, 45). To investigate this possibility, we performed immunostaining for PECAM-1, an endothelial marker associated with cell–cell junction integrity, and vimentin, a marker indicative of enhanced cell motility.

Under flow conditions, the nuclei were aligned along the channel walls, and PECAM-1 was localized at cell–cell junctions, while vimentin expression remained weak. This combination resulted in the formation of a well-defined vascular lumen (Fig. 6C, D, F). This cellular morphology closely resembles that of typical blood vessels, indicating the successful formation of physiological blood vessels under flow conditions. These findings suggest that the maintenance of cell–cell junctions and collective cell migration are essential for effective vascular formation within the channels.

In contrast, under static conditions where vessel formation failed, nuclear alignment was disrupted, and PECAM-1 was not localized at cell–cell junctions but instead exhibited uniform distribution across the cell membrane (Fig. 6E, G). In these regions, vimentin expression was elevated, consistent with the high degree of individual cell motility observed in time-lapse imaging (Fig. 5C). These findings suggest a loss of cell polarity, increased cell motility, and reduced cell–cell adhesion under static conditions, indicative of EndMT (44, 46, 48). The inability of endothelial cells to form a monolayer along the channels under static conditions can likely be attributed to the loss of apical-basal polarity caused by EndMT, as cells adhered to the channel walls in all directions (Fig. 4E).

The establishment of apical-basal polarity depended on the size of channels relative to the size of the cells. In channels with a diameter of 50 µm, which is significantly larger than the size of individual cells, the cells adhered to one side of the luminal surface, resembling their behavior in 2D culture. Consequently, the cell–ECM interface formed the basal domain, while the cell surface exposed to the culture medium became the apical domain, thereby establishing apical-basal polarity. In contrast, in channels with a diameter of 20 µm, some cells adhered to the luminal surface in all directions, disrupting the formation of a distinct basal domain and preventing the establishment of apical-basal polarity. Without the establishment of apical-basal polarity, cells cannot form a lumen (49). The application of flow may help align cells along the luminal surface, thereby promoting the formation of a basal domain and establishing an apical domain oriented toward the culture medium.

### Engineering predefined microvascular architecture and its applications

In this study, we established a method to fabricate hollow channels with sizes in the range of capillaries and microvessels, featuring any desired shape within hydrogels, such as collagen gels and fibrin-collagen gels. One of the key advantages of this method is its ability to reproducibly generate microvascular networks *in vitro* with predefined shapes. In previous studies, controlling the shape of microvascular networks has been challenging, as it relied on the spontaneous formation of networks by endothelial cells. In contrast, our method enables the reproducible creation of vascular networks with precisely defined geometries.

We employed a simple straight geometry to investigate the key factors required for vascular formation in the range of capillaries and microvessels. This approach enabled highly reproducible experiments under both static and flow conditions, providing insights into the vascular formation processes. Additionally, this experimental system facilitated detailed observations of cellular polarization, migration, and other morphological changes during vascular formation. Beyond straight vasculature, our system supports the formation of branched and looped vascular networks (Fig. 7), enabling detailed analysis of remodeling processes such as vessel pruning. In this study, we demonstrated the expansion of the diameter, which is one of the typical remodeling of microvessels.

The reproducibility offered by this method holds significant potential for constructing organ-specific vascular networks in tissue engineering. Organs such as the liver and kidney have unique vascular architectures tailored to their metabolic demands, making the replication of such specific 3D vascular arrangements crucial for successful tissue regeneration. A pioneering work has demonstrated the construction of vascular networks mimicking kidney glomeruli (36). To enhance physiological relevance, future studies using this model should also consider co-culture with perivascular cells such as pericytes and fibroblasts. Additionally, this system can be applied to epithelial cells to construct organ-specific epithelial tissues, such as those of the liver and kidney. Integrating microvascular networks with parenchymal cells presents a promising approach for generating vascularized 3D architectures in predefined shapes *in vitro*.

The limitations of this model are its low throughput and limited scalability. First, each channel requires approximately 30 minutes to fabricate using femtosecond laser ablation with our current system, indicating the need for improvements in processing speed. Second, the scale at which ablation can be performed is constrained by the working distance of the objective lens, making it challenging to fabricate structures over organ-scale areas spanning several tens of centimeters. However, it is possible to fabricate modular structural units using this method and assemble them to construct larger tissue architectures.

## Concluding remarks

This study presents a method for constructing vascular networks with predefined shapes using multiphoton lithography, specifically targeting the capillary and microvessel size range. We identified key differences in vascular formation under flow and static conditions at the capillary scale, highlighting the critical roles of cell–cell adhesion and motility. These findings on the vascular formation along microchannels represent a significant advancement in engineering organ-specific microvascular networks *in vitro*. Additionally, our culture model, which provides controlled vascular geometry and a stable flow environment, serves as a valuable tool for investigating vascular responses to various stimuli, including vascular remodeling under both physiological and pathological conditions. More importantly, this model holds great potential for in-depth studies of engineering complex 3D architectures that integrate hierarchical vascular networks and epithelial tissues *in vitro*, particularly in achieving predefined geometries.

## Abbreviations

2D: two dimensional
3D: three dimensional
EndMT: endothelial-to-mesenchymal transition
PDMS: polydimethylsiloxane
HUVEC: human umbilical vein endothelial cell
EGM-2: endothelial growth medium-2
PBS: phosphate buffered saline
PECAM-1: platelet endothelial cell adhesion molecule-1
PIV: particle imaging velocimetry
ECM: extracellular matrix

## Acknowledgements

This study was supported by the Japan Society for the Promotion of Science (grant number: 19H04452, 23H04089, 25K03439), Keio Leading-edge Laboratory of Science and Technology, and JST SPRING (grant number: JPMJSP2123).

## Author Contributions

A.O. and R.O. designed and conducted experiments, collected, and analyzed the data. Y.K., M.S. and Y.T. designed and performed the multiphoton ablation experiments. T.Y. interpreted experimental results and conducted image correlation analysis of time-lapse movies. R.S. conceived and designed the study, supervised experiments, and interpreted experimental results. A.O. wrote the original draft, and R.S. reviewed and revised the manuscript. All authors edited and approved the final manuscript.

---

**Supplementary Fig. 1.**
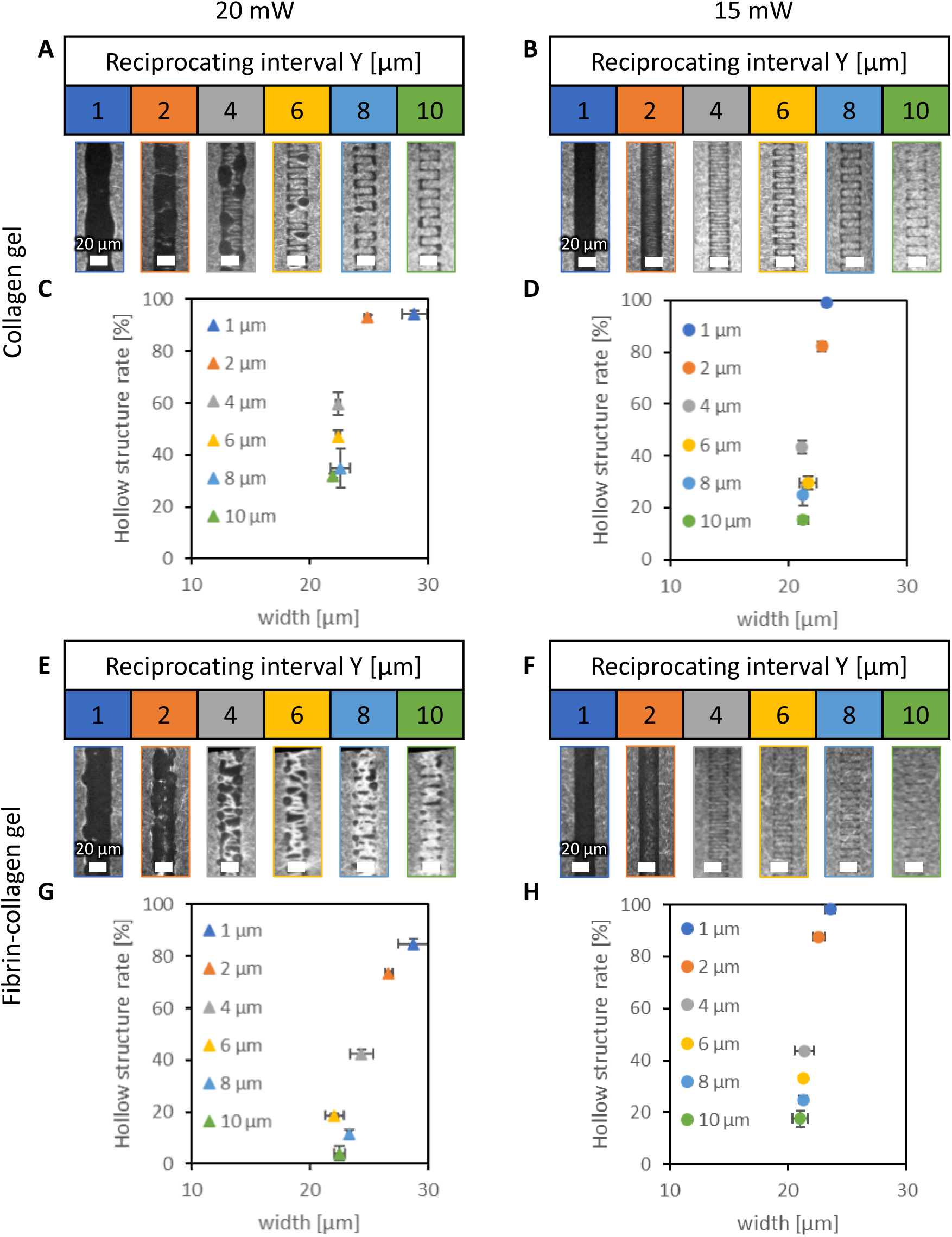
Hollow structures fabricated along single-layer zigzag trajectory at various reciprocating intervals. (A, B) Confocal reflection images of the hollow structures in collagen gel fabricated along the single-layer trajectory at irradiation intensities of 20 and 15 mW. Scale bars, 20 μm. (C, D) Quantification of the width of the hollow structures in collagen gel. (E, F) Confocal reflection images of the hollow structures in fibrin-collagen gel fabricated along the single-layer trajectory at irradiation intensities of 20 and 15 mW. Scale bars, 20 μm. (G, H) Quantification of the width of the hollow structures in fibrin-collagen gel. Data are shown as the mean ± SD. n=3.

**Supplementary Fig. 2.**
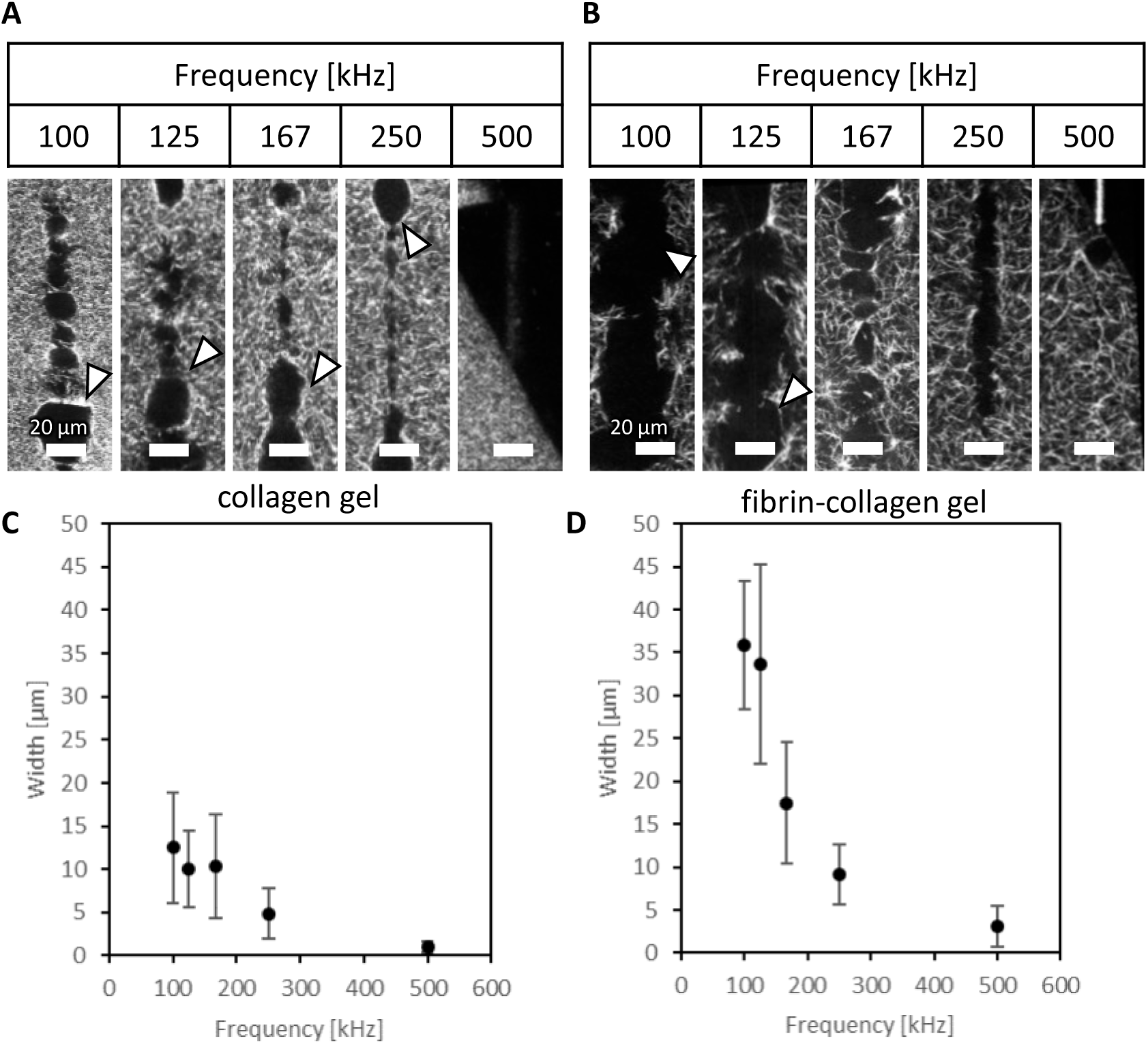
Hollow structures fabricated along single-layer straight-line trajectory at various frequencies. (A, B) Confocal reflection images of the hollow structures in collagen gel and fibrin-collagen gel fabricated along the single-layer straight-line trajectory. Scale bars, 20 μm. (C, D) Quantification of the width of the hollow structures in collagen gel and fibrin-collagen gel. Data are shown as the mean ± SD. n=51.

**Supplementary Fig. 3.**
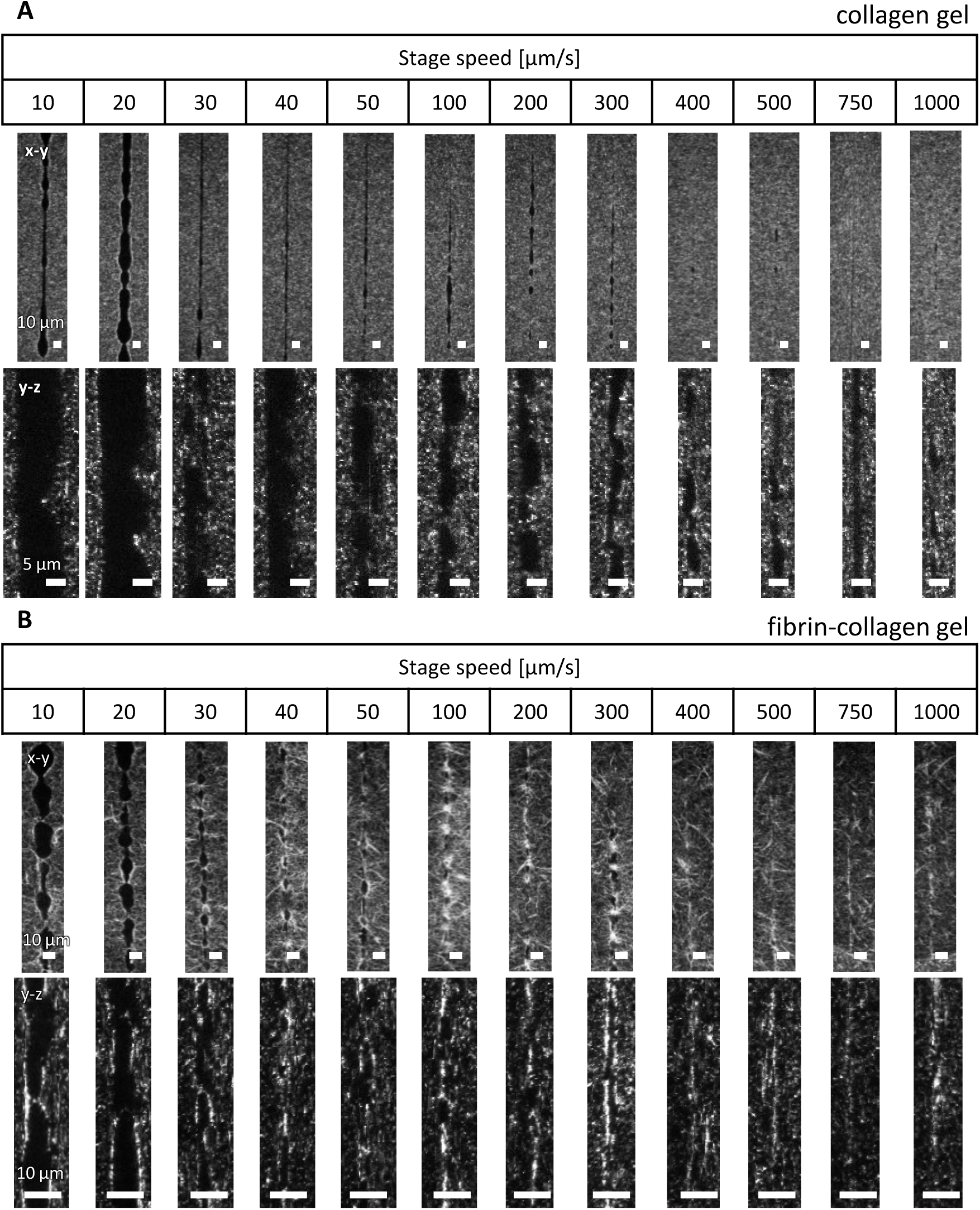
Hollow structures fabricated along single-layer straight-line trajectory at various stage speed. (A) Confocal reflection images of the hollow structures in collagen gel fabricated along the single-layer straight-line trajectory. Scale bars, 10 μm. (B) Confocal reflection images of the hollow structures in fibrin-collagen gel fabricated along the single-layer straight-line trajectory. Scale bars, 10 μm.

**Supplementary Fig. 4.**
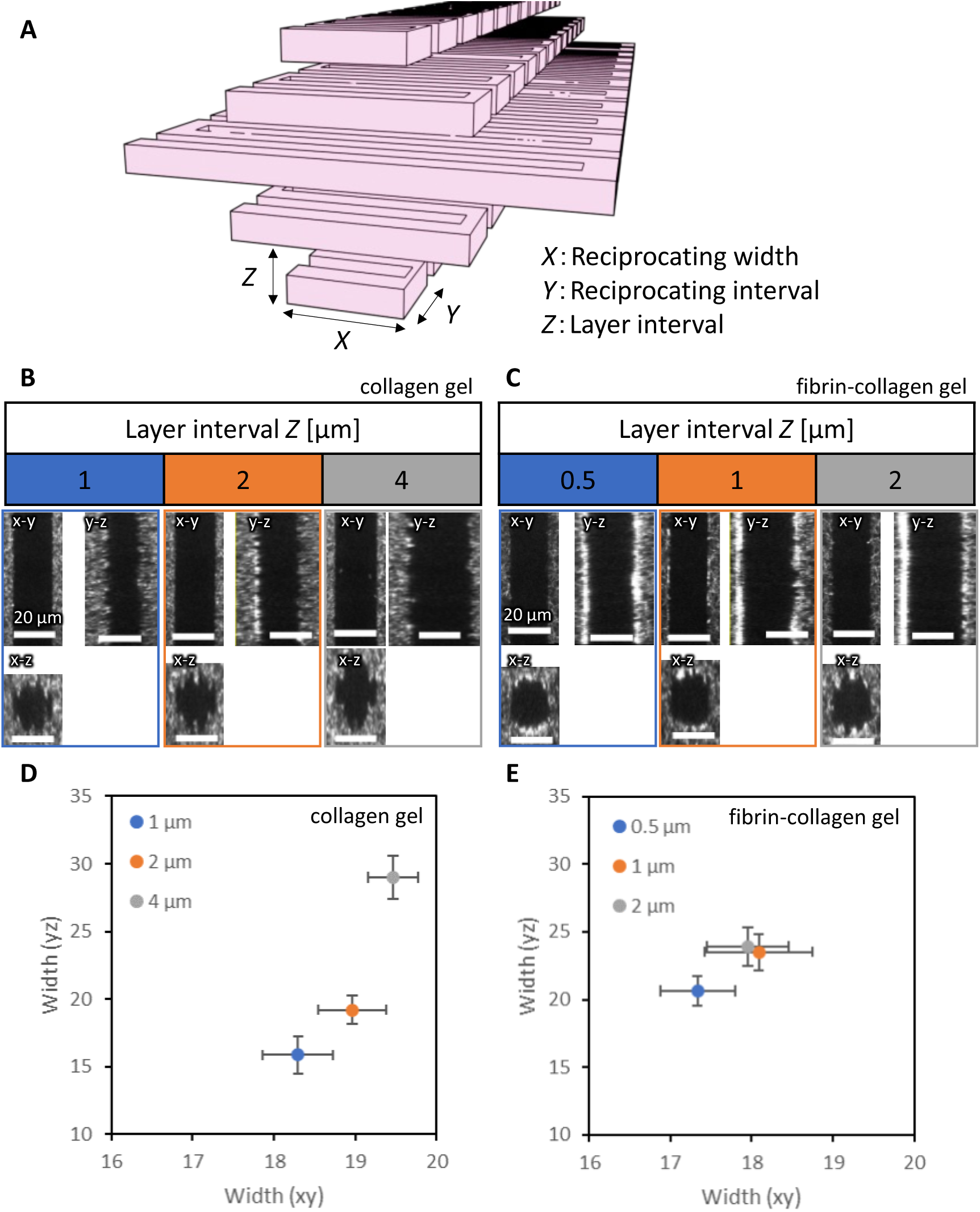
Hollow structures fabricated along five-layer zigzag trajectory at various irradiation intervals. (A) Schematic illustration of the five-layer zigzag trajectory. (B, C) Confocal reflection images of the hollow structures in collagen gel and fibrin-collagen gel fabricated along the five-layer zigzag trajectory. Scale bars, 20 μm. (D, E) Quantification of the width of the hollow structures in collagen gel and fibrin-collagen gel. Data are shown as the mean ± SD. n=51.

**Supplementary Fig. 5.**
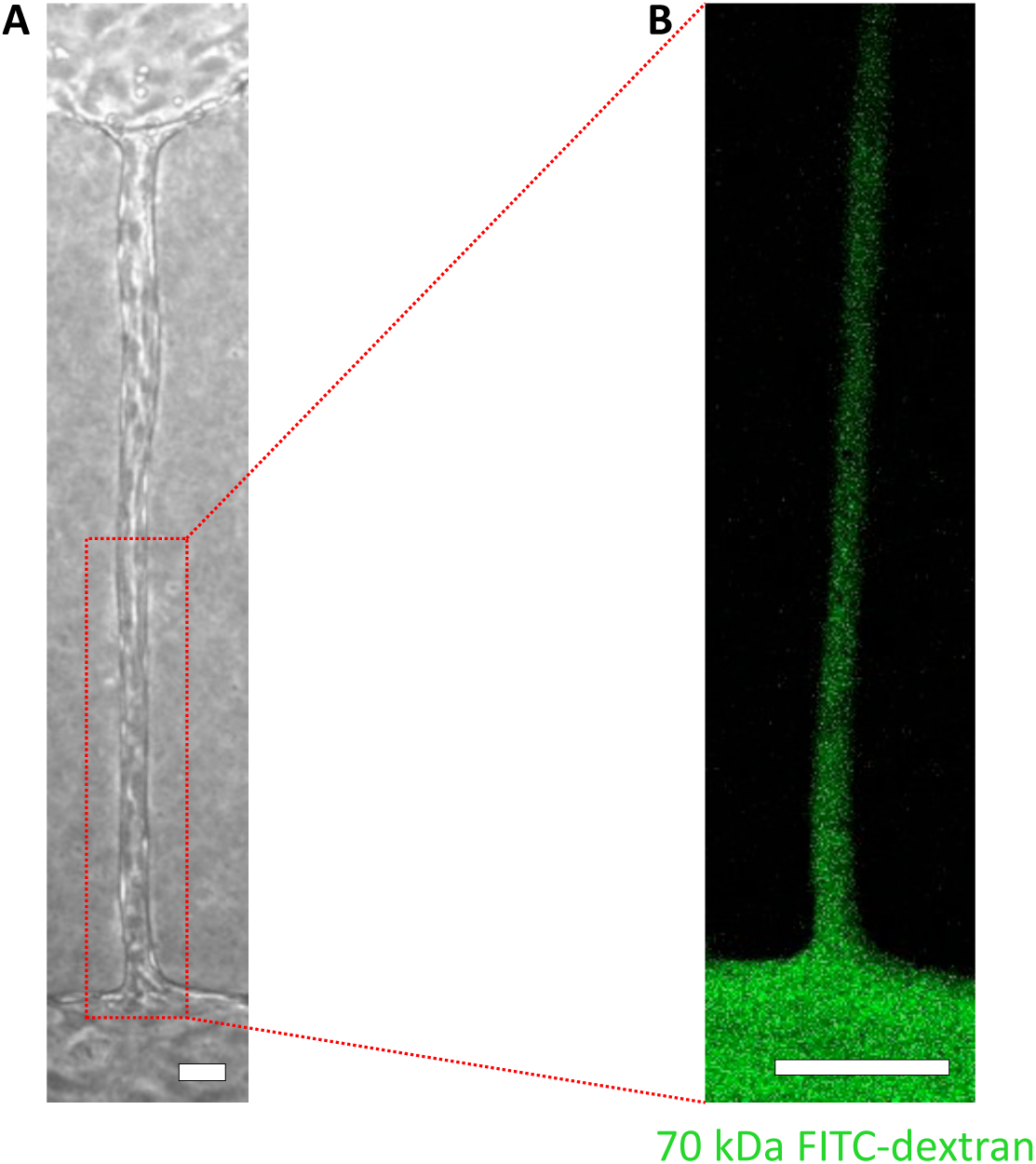
Perfusability and endothelial barrier function of microvessels formed along 20 µm microchannels. (A) Phase-contrast image of a microvessel formed under flow conditions. Scale bar, 50 μm. (B) Confocal fluorescence images showing perfusion of 70 kDa FITC-dextran through the microvessel. Scale bar, 100 μm.

**Supplementary Fig. 6.**
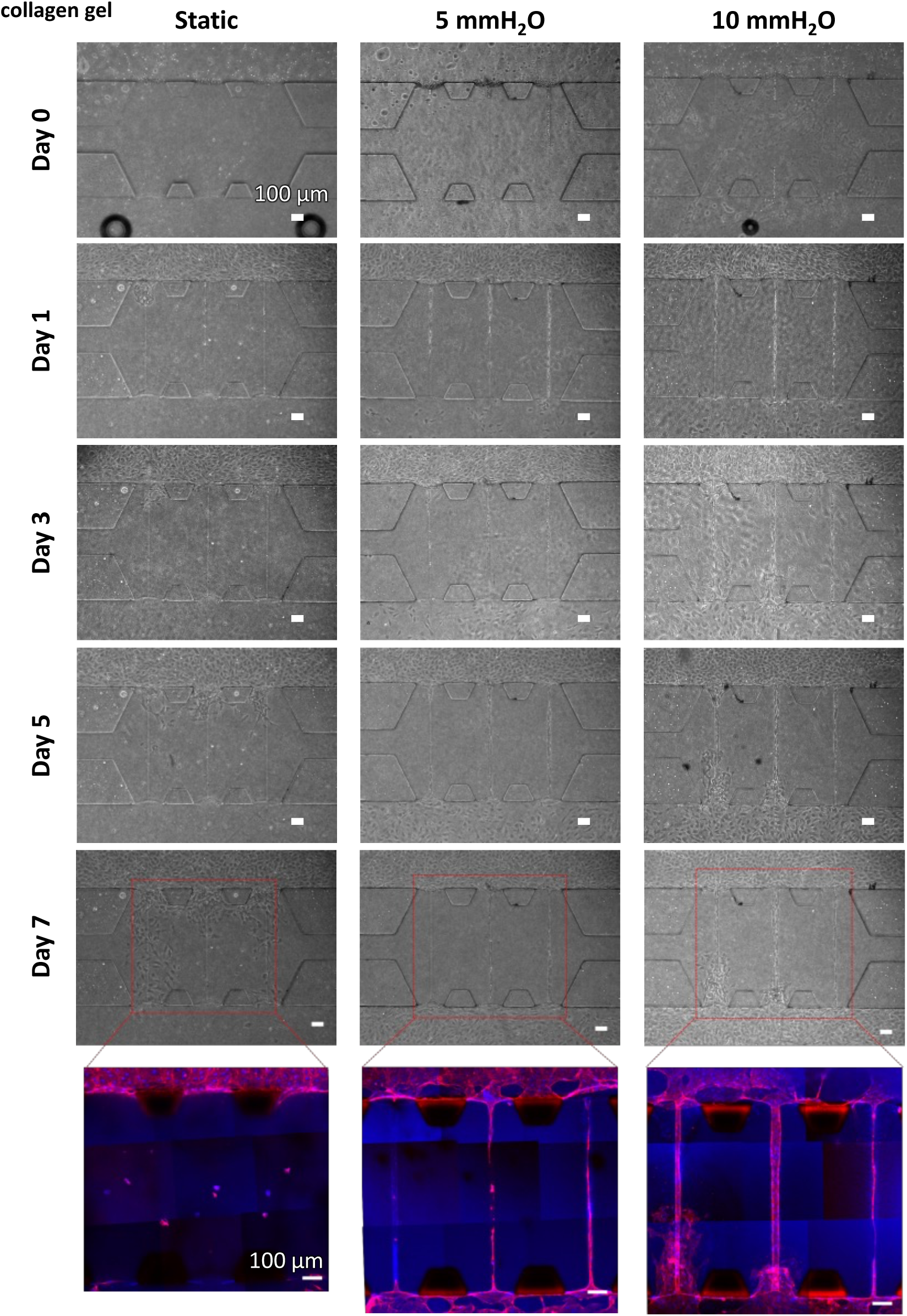
Vascular formation along a 20 μm diameter channel in collagen gel. (A) Phase-contrast images of vascular formation under static and flow condition (0, 5, 10 mm H_2_O). Scale bars, 100 μm. (B) Immunofluorescence images of vascular formation under static and flow condition (0, 5, 10 mm H_2_O). Cells were fixed on day 7 and stained for nuclei (Hoechst 33342) and F-actin (phalloidin). Scale bars, 100 μm.

**Supplementary Fig. 7.**
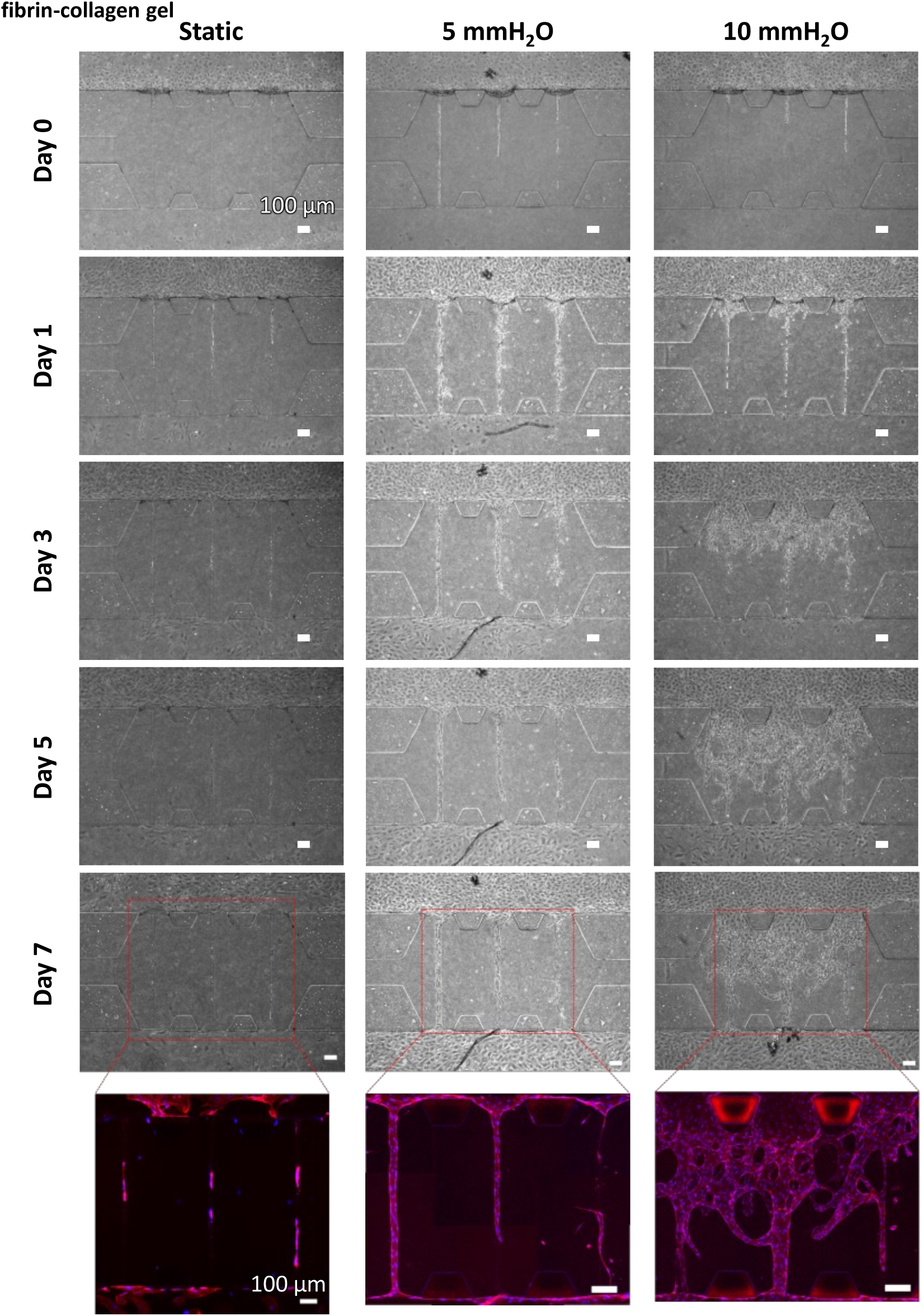
Vascular formation along a 20 μm diameter channel in fibrin-collagen gel. (A) Phase-contrast images of vascular formation under static and flow condition (0, 5, 10 mm H_2_O). Scale bars, 100 μm. (B) Immunofluorescence images of vascular formation under static and flow condition (0, 5, 10 mm H_2_O). Cells were fixed on day 7 and stained for nuclei (Hoechst 33342) and F-actin (phalloidin).

**Supplementary Movie 1 Time-lapse movie of the cells under static conditions.** Sequential phase-contrast images were obtained by phase-contrast microscope equipped with a time-lapse imaging system. Images were obtained at 15 min intervals for 24 h starting on day 2.

**Supplementary Movie 2 Time-lapse movie of the cells under flow conditions (10 mmH_2_O).** Sequential phase-contrast images were obtained by phase-contrast microscope equipped with a time-lapse imaging system. Images were obtained at 15 min intervals for 24 h starting on day 2.

**Supplementary Movie 3 Image correlation analysis movie under static conditions.** Supplementary movie 1 was analyzed for visualizing the movement of each cell.

**Supplementary Movie 4 Image correlation analysis movie under flow conditions.** Supplementary movie 2 was analyzed for visualizing the movement of each cell.

